# Multimodal gradients of basal forebrain connectivity across the neocortex

**DOI:** 10.1101/2023.05.26.541324

**Authors:** Sudesna Chakraborty, Roy A.M. Haast, Kate M. Onuska, Prabesh Kanel, Marco A.M. Prado, Vania F. Prado, Ali R. Khan, Taylor W. Schmitz

## Abstract

The cholinergic innervation of the cortex originates almost entirely from populations of neurons in the basal forebrain (BF). Structurally, the ascending BF cholinergic projections are highly branched, with individual cells targeting multiple different cortical regions. However, it is not known whether the structural organization of basal forebrain projections reflects their functional integration with the cortex. We therefore used high-resolution 7T diffusion and resting state functional MRI in humans to examine multimodal gradients of BF cholinergic connectivity with the cortex. Moving from anteromedial to posterolateral BF, we observed reduced tethering between structural and functional connectivity gradients, with the most pronounced dissimilarity localized in the nucleus basalis of Meynert (NbM). The cortical expression of this structure-function gradient revealed progressively weaker tethering moving from unimodal to transmodal cortex, with the lowest tethering in midcingulo-insular cortex. We used human [^18^F] fluoroethoxy-benzovesamicol (FEOBV) PET to demonstrate that cortical areas with higher concentrations of cholinergic innervation tend to exhibit lower tethering between BF structural and functional connectivity, suggesting a pattern of increasingly diffuse axonal arborization. Anterograde viral tracing of cholinergic projections and [^18^F] FEOBV PET in mice confirmed a gradient of axonal arborization across individual BF cholinergic neurons. Like humans, cholinergic neurons with the highest arborization project to cingulo-insular areas of the mouse isocortex. Altogether, our findings reveal that BF cholinergic neurons vary in their branch complexity, with certain subpopulations exhibiting greater modularity and others greater diffusivity in the functional integration of their cortical targets.

## Introduction

The basal forebrain (BF) (Fig. 1) is a collection of subcortical cholinergic cell groups which provide the major sources of acetylcholine to the cortex and hippocampus^1^. Structurally, the ascending cholinergic projections are highly branched, with individual cells often targeting multiple different cortical areas^2–4^. The total arborization of a single human cholinergic BF neuron is estimated to have a length in excess of 100 meters^4^.

**Fig. 1.**
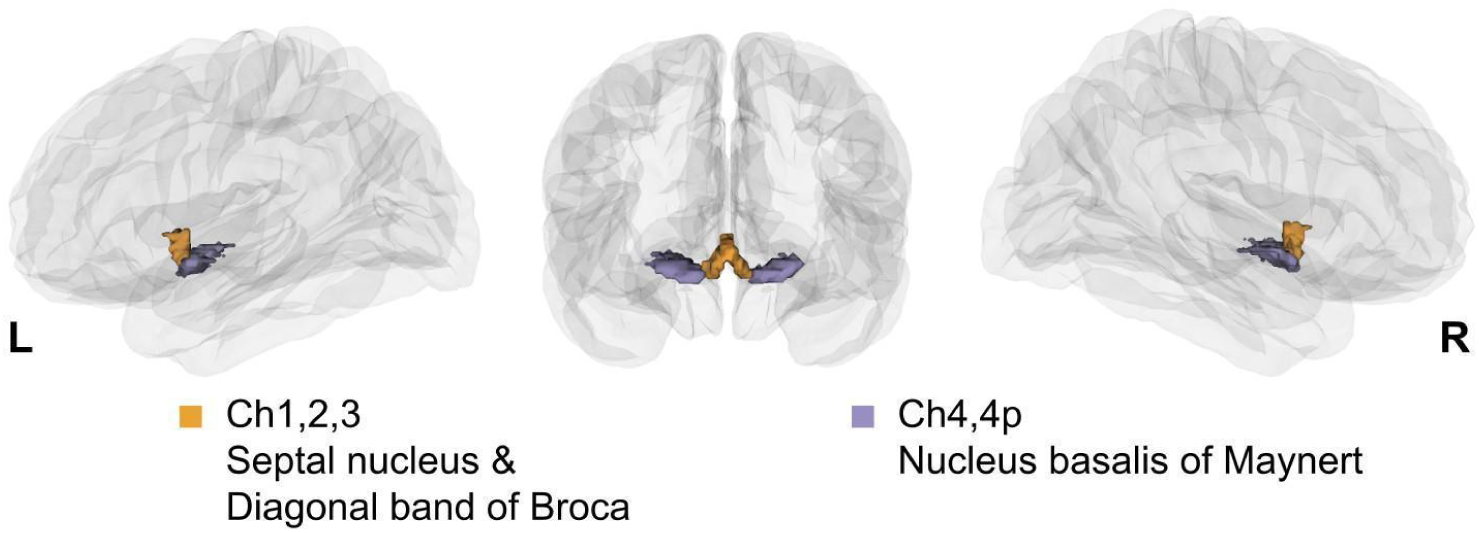
A 3D view of histologically defined BF subdivisions defined by Zaborsky et al^28^ projected on a glass brain. The anteromedial nuclei are displayed in yellow and posterolateral nuclei in purple.

The organization of ascending BF cholinergic projections may reflect complex spatial topographies of connectivity with the cortex^5,6^. Axonal tracing studies in rats suggest that BF cholinergic neurons are grouped into ensembles which target functionally interrelated cortical areas^7^. In humans, patterns of functional connectivity in distinct BF subregions have been found to overlap with distinct cortico-cortical networks^8–10^. Consistent with a spatial topography of projections, subregional structural changes in BF gray matter and white matter integrity are associated with distinct patterns of cortical degeneration and cognitive dysfunction^11–15^. In neurodegenerative diseases such as Alzheimer’s (AD), early dysfunction or loss of specific BF cholinergic fibers may alter local neuronal functions in cholinoreceptive cortical areas^16^. Although these separate lines of evidence suggest that the cortex expresses topographies of BF structural and functional connectivity, the interrelationship of these topographies to one another is unknown.

At the cortico-cortical level, the degree to which interregional structural connectivity can predict their functional integration, also known as structural-functional coupling or ‘tethering’, appears to decrease from unimodal to transmodal areas and to coincide with functional and cytoarchitectonic cortical hierarchies^17–20^. However, how does the structural organization of ascending cholinergic BF projections relate to their functional integration in the cortex? One possibility is that BF structural and functional connectivity are closely tethered. This would be expected if white matter tracts emanating from local populations of BF cholinergic neurons target local patches of cortex^5^. In this scenario, BF functional connectivity would reflect these structurally segregated cortical modules. Alternatively, BF structural and functional connectivity might exhibit negligible overlap. This pattern would be expected if the BF white matter tracts are highly diffuse, with each local population of cholinergic neurons projecting to multiple distinct and overlapping cortical areas^5^. In this scenario, BF functional connectivity would reflect mixtures of these diffuse afferent inputs. Finally, a third possibility is that the profile of tethering between BF structural and functional connectivity varies depending on the properties of different cortical areas^18,21^ and the biophysical constraints these properties may impose on the cholinergic projections^22,23^ . This pattern would be expected if populations of BF cholinergic neurons vary in their branch complexity, with certain subpopulations exhibiting greater modularity and others greater diffusivity in the functional integration of their cortical targets.

Here we addressed the relationship between structural and functional connectivity in the ascending BF projections. We used multimodal imaging combining high-resolution 7 Tesla (7T) diffusion (dMRI) and resting state functional MRI (rsfMRI) in a cohort of 173 individuals from the Human Connectome project (HCP)^24^. From these datasets (Supplemental Table 1), we derived gradients of the continuous transitions in BF functional and structural connectivity with the cortex using diffusion map embedding^25^. Diffusion map embedding is a statistical technique that has recently gained traction to extract different topographies or ‘gradients’ of the continuous transitions in structural and functional connectivity within the brain (i.e., entire cortex or individual regions^26,27^). In both the dMRI and rsfMRI datasets, a single gradient dominated the explained variance in BF connectivity, characterized by an anteromedial to posterolateral axis spanning the BF nuclei. To quantify how these patterns of BF structural and functional connectivity were tethered, we computed the residual value between (a) the gradients of structural and functional connectivity within the BF and (b) the expression of these gradients on the cortical surface. Consistent with a gradient of branch complexity across the BF, we found closer tethering (lower residual values) between BF structural and functional connectivity in anteromedial areas of the BF, with stronger divergence (higher residual values) in posterolateral areas overlapping the nucleus basalis of Meynert (NbM). The cortical expression of BF structure-function tethering itself exhibited a gradient, with the closest tethering observed in primary sensory cortices and greatest divergence in midcingulo-insular hubs of the ventral attention network^28–36^. Using *in vivo* molecular imaging of the human^37,38^ and mouse cortical cholinergic projections, in combination with anterograde viral labeling of individual BF cholinergic axons in mice (Supplemental Table 1)^2^, we provide confirmatory cell-type specific evidence that the observed gradient of tethering between BF structural and functional connectivity is likely shaped by the complexity of axonal arborization in BF cholinergic neurons. In mouse and human, BF cholinergic neurons with greater arborization appear to target evolutionarily conserved midcingulo-insular cortical areas which exhibit among the highest concentrations of cortical cholinergic innervation.

## Results

### Primary BF structural and functional gradients

We built dMRI and rsfMRI connectomes of the BF using high-resolution 7T MRI HCP data (*n*=173; Supplemental Table 1)^39^ and a well-validated stereotactic BF atlas which provides annotations for its separate nuclei (Fig. 1)^40^. For each individual, the structural and functional connectomes were separately defined by an *M*-by-*N* matrix, where *M* represents the voxels in the entire BF and *N* represents the cortical parcels of the HCP multimodal parcellation (HCP-MMP)^41^. For rsfMRI data, the matrices encode the interregional temporal correlations in blood oxygenation level-dependent signals. For dMRI data, the matrices encode the streamline counts between different regions of the brain and BF. We confirmed that the probabilistic tractography of the BF streamlines recapitulates prior ex vivo anatomical tracing^42^ and in vivo diffusion mapping^14,43^ work on the human BF projection system which distinguish midline cingulum and lateral capsular/perisylvian white matter projection routes (Supplemental Fig. 1).

For the dMRI and rsfMRI datasets, we computed gradients of connectivity using the BrainSpace toolbox^44^. Starting from the input matrices (Supplemental Fig. 2A) encoding the rsfMRI and dMRI data (averaged across individuals), we used a kernel function to build an affinity matrix encoding interregional similarity of features (Supplemental Fig. 2B). These matrices were decomposed via nonlinear dimension reduction into a set of eigenvectors describing axes of BF connectivity with cortical targets. The axes were then projected back to the BF space (Supplemental Fig. 2C).

The first gradient for both BF structural and functional connectivity data explained the most (30%) variance, with a drop to 15% explained variance for the second gradient (Fig. 2A). We therefore focused on the first principal gradient, which exhibited a smooth continuous transition from anteromedial to posterolateral BF for both the structural and functional connectomes (Fig. 2B,C). The connectivity profiles are consistent with findings from histological mapping of the individual BF nuclei^12,40^ and in prior cluster-based parcellations of the BF from rsfMRI^8–10^. Note that the hemispheric asymmetry in these gradients is due to the original stereotaxic mapping of the BF atlas (Fig. 2B; Methods).

**Fig. 2.**
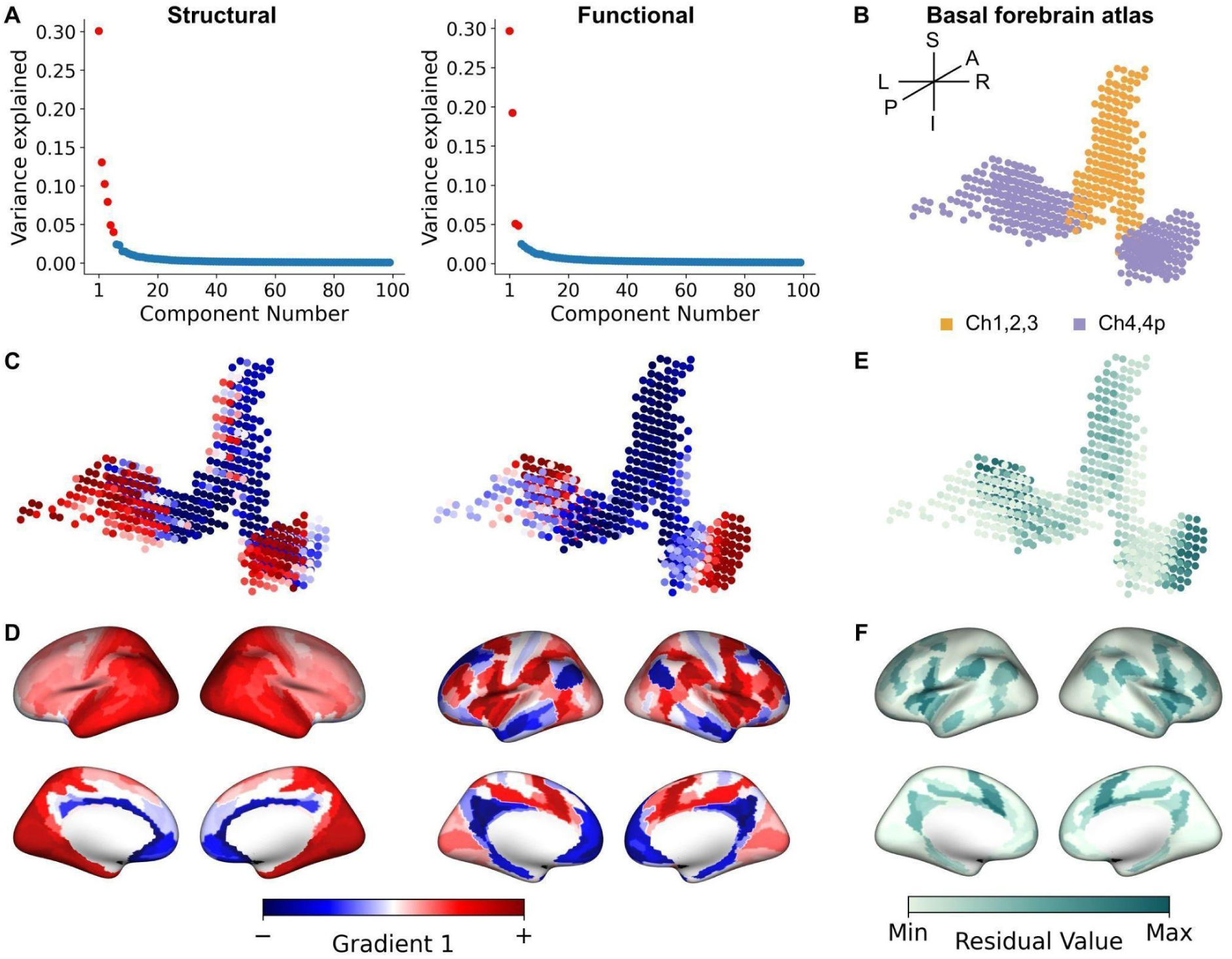
Basal forebrain (BF) gradients of structural and functional connectivity. (A) Scree plots showing the variance explained by each gradient of structural (left) and functional connectivity (right). Gradients falling above the knee-point^34^ are denoted as red. (B) The BF atlas projected into 599 voxels color coded according to a priori histologically defined anteromedial or posterolateral nuclei. (C) The first principal gradient of the BF based on structural (sG1; left) and functional (fG1; right) connectivity both revealed an anteromedial to posterolateral axis. The lower bound of gradient values is represented by blue while the upper bound is represented by red. (D) Gradient-weighted cortical maps corresponding to sG1 (left) and fG1 (right). (E) Residual values encoding tethering between structural and functional connectivity at each BF voxel. Darker green values indicate lower tethering between structural and functional connectivity. (F) Gradient-weighted cortical maps corresponding to structure-function tethering.

We next computed gradient-weighted cortical maps^45^ to determine how BF gradients were expressed by the cortex. The gradient-weighted cortical maps were created by multiplying each row of the initial connectivity matrix (*M*_BF voxels_ x *N*_cortical parcels_) with the corresponding structural principal gradient (sG1) or functional principal gradient (fG1) value to create a gradient-weighted connectivity matrix. Finally, all rows of this gradient-weighted matrix were averaged to produce a single cortical representation of the particular gradient (see Supplemental Fig. 2E and Methods). For the gradient-weighted cortical map corresponding to sG1 (sG1ctx; Fig. 2D left), we observed a smooth macroscale transition from the anteromedial to posterolateral cortical surface. By contrast, the gradient-weighted cortical map corresponding to fG1 (fG1ctx; Fig. 2D right) exhibited a more patch-like pattern with moderate differentiation of gradient weights between midline and lateral cortical areas.

### A gradient of structure-function tethering in BF connectivity

We examined the magnitude of shared variance, or tethering, between BF structural and functional gradients to determine their spatial similarity to one another. To do so, we computed voxel-wise regressions for the fG1 and sG1 structure-function gradient pair and extracted their corresponding residuals (see Methods). The residual map encoding BF structure-function tethering was projected back to BF space (Supplemental Fig. 2D), which revealed an anteromedial to posterolateral topography, with the lowest structure-function tethering localized in posterolateral subregions (Fig. 2E). We next computed the gradient-weighted cortical map encoding our residualized map of BF structure-function tethering. The resulting cortical map (Fig. 2F) exhibited decreased tethering moving from unimodal to transmodal cortex, with lowest tethering in the anterior cingulate, insular and frontal opercular cortices.

How does the smooth transition of tethering between BF structural and functional connectivity (Fig. 2E) align with the histologically defined boundaries of the BF anteromedial (Ch1,2,3) and posterolateral (Ch4,4p) nuclei (Fig. 2B)^40^? To address this question, we examined if the distributions of residual values within Ch1,2,3 and Ch4,4p differed from one another. We performed permutation tests with fitted surrogate maps (see Methods) to compute the difference in both the means and dispersion between Ch1,2,3 and Ch4,4p. Dispersion was quantified by the coefficient of variation (CoV) defined as the ratio of the standard deviation to the mean of the data. The mean residual values were not different between Ch1,2,3 and Ch4,4p (p_perm_=0.40). However, the residual values in Ch4,4p exhibited significantly greater variability in comparison to Ch1,2,3 (p_perm_=0.01). This latter finding suggests that structure-function tethering within Ch4,4p was more inhomogeneous than Ch1,2,3 (Fig. 3A).

**Fig. 3.**
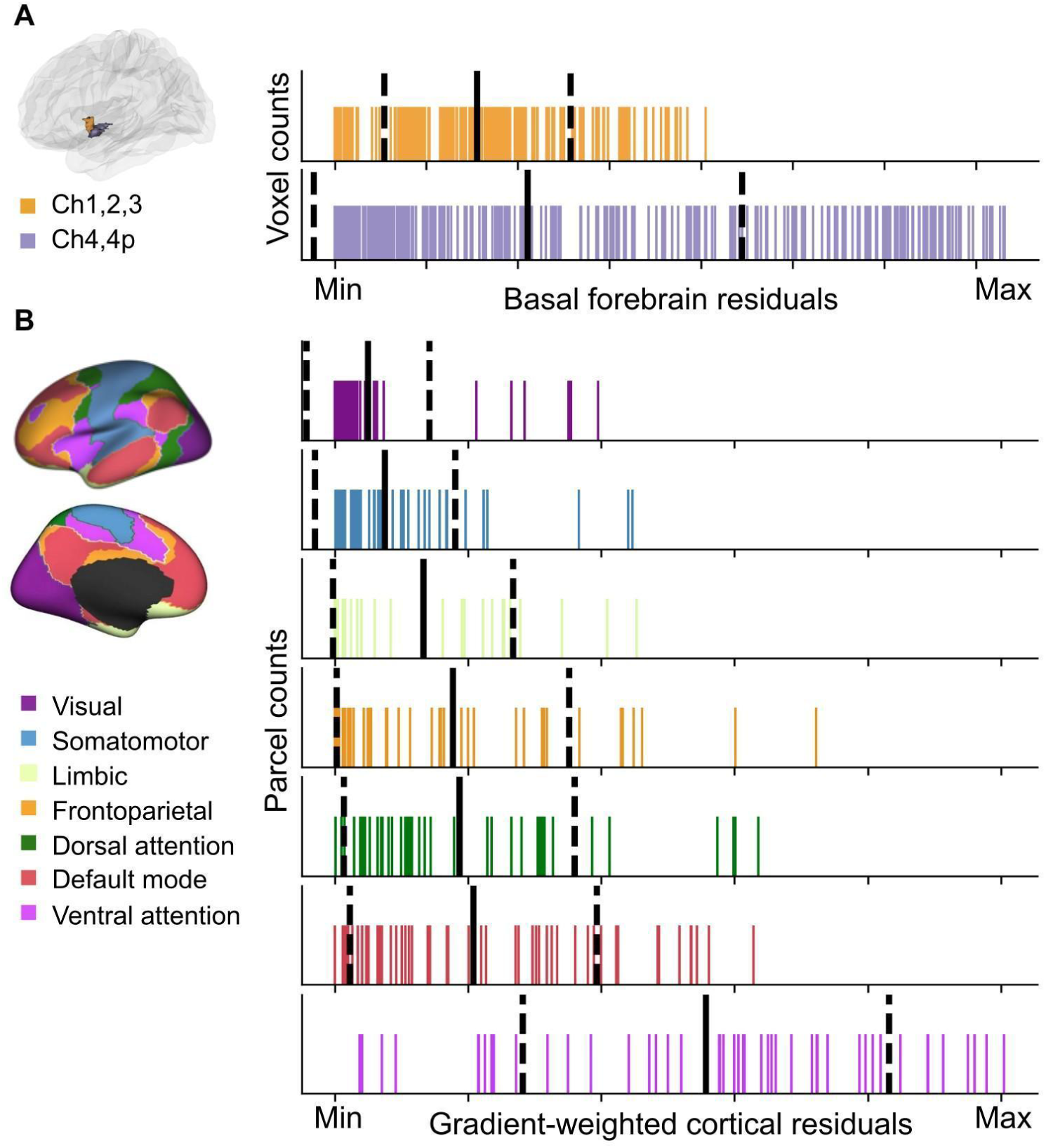
Distributions of basal forebrain (BF) and cortical residuals. (A) Rug plot showing the distribution of residuals within the *a priori* histologically defined anteromedial (Ch1,2,3) and posterolateral (Ch4,4p) BF nuclei. The solid black line indicates the mean and the dashed lines indicate +/-one standard deviation. The minimum and maximum values correspond to the same scale as in Fig 2E. (B) Rug plot showing the distribution of residuals within each of the 7 intrinsic resting state networks identified by Yeo et al^25^. The color-coding for different networks is based on the original parcellation. The solid black line indicates the mean and the dashed lines indicate +/-one standard deviation. The minimum and maximum values correspond to the same scale as in Fig 2F.

We then asked how well the cortical residual-weighted map of BF structure-function tethering (Fig. 2F) aligns with the spatial topographies of intrinsic cortico-cortical resting state networks. To do so, we compared the mean and CoV of the distributions of residual values captured by each of 7 Yeo macroscale resting state networks^35^ covering the entire human cerebral cortex. A pattern in which the distributions of these residual values are well delineated from one another across the different cortico-cortical networks (distinct means, lower CoVs) would indicate that each network falls along a continuum of BF structure-function tethering defined by the observed anteromedial-to-posterolateral gradient. By contrast, a pattern in which the distributions of residuals are more spread out across the different cortico-cortical networks (overlapping means, higher CoVs) would indicate that each network exhibits a mixture of lower and higher BF structure-function tethering. We found that the mean of the residuals significantly differed among networks (F_6,330_ =47.55, p=0.001, Fig. 3B), with the ventral attention network characterized by significantly higher values than the visual (p_perm_=0.001) and somatomotor networks (p_perm_=0.001). The dorsal attention, limbic, frontoparietal and default mode networks exhibited an intermediate tendency for structure-function tethering. However, we also observed that the CoVs captured by each cortico-cortical network also differed. Again, the ventral attention network exhibited significantly greater heterogeneity in residual values than the visual (p_perm_=0.001) and somatomotor networks (p_perm_=0.001; Fig. 3B). Taken together, these findings support a continuum of BF structure-function tethering. Cortico-cortical networks with hubs primarily in the unimodal cortex (visual, somatomotor) tend to exhibit higher tethering, while networks with hubs primarily in the transmodal cortex tend to exhibit lower tethering.

To assess the stability of the dominant structural and functional connectivity gradients (sG1 and fG1) and the gradient of their tethering, we performed cross-validation analyses across the 173 individuals in the sample (see Methods). We found that each individual’s observed gradients were consistently highly correlated with the model (split-half or leave-one-out), consistent with stable gradient organization across individuals (Supplemental Fig. 3).

Having confirmed the stability of sG1, fG1, and their tethering across individuals, we next examined if gradients with lower explained variance, i.e. sG2 and fG2, may reveal additional profiles of BF structure-function tethering. To explore this possibility, we computed the ‘knee-point’ in the curve of explained variance for the complete set of observed structural and functional gradients (see Methods), where the change in explained variance became negligible^46^. The knee-point corresponded to sG6 for structural gradients and fG4 for functional gradients (Fig 2A, red versus blue color gradient component). For each of the 24 structure-function gradient pairs in this 6 x 4 matrix (Supplemental Fig. 4A), we computed the structure-function tethering metric as before. When we examined the average of the 24 residual maps (Supplemental Fig. 4B), we observed a high degree of correspondence to the sG1-fG1 residual map (r=0.918; Supplemental Fig. 4C), even when excluding sG1-fG1 pairing from the average (r=0.878; Supplemental Fig. 4D). These findings imply that the gradient of tethering between BF structural and functional connectivity is captured by a single dominant profile (Fig. 2E).

### BF structure-function tethering reflects an arborization gradient of cholinergic neurons

We next explored what properties of BF connectivity might shape the observed gradient profile of structure-function tethering in the cortex. Currently, [^18^F]FEOBV positron emission tomography (PET) provides the most direct *in vivo* assay of cortical cholinergic innervation in humans. The [^18^F]FEOBV radiotracer binds to the vesicular acetylcholine transporter (VAChT), a glycoprotein expressed solely by cholinergic neurons, with the highest density of expression on the presynaptic terminals^47^. We acquired distribution volume ratios (DVR) of [^18^F]FEOBV binding from a group of healthy cognitively normal young adults (*n*=13; mean age=24.54, 3 females, Supplemental Table 1)^38^, and produced an average map representing BF cortical cholinergic innervation (Fig. 4A; see Methods). We then examined the correlation between the cortical expression of BF structure-function detethering (cortical residual map from Fig. 2F) and cholinergic innervation estimated from the average [^18^F]FEOBV map using spatial spin tests^48^. Surprisingly, we found that cortical areas exhibiting lower BF structure-function tethering, i.e., more branching, exhibited a higher density of BF cholinergic input, i.e., higher VAChT concentration (R=0.28, p_spin_=0.02; Fig. 4B). We replicated these associations with three other publicly available atlases of [^18^F]FEOBV PET^37,49,50^ (Supplemental Table 1) and found similar strong positive relationships (Supplemental Fig. 5).

**Fig. 4.**
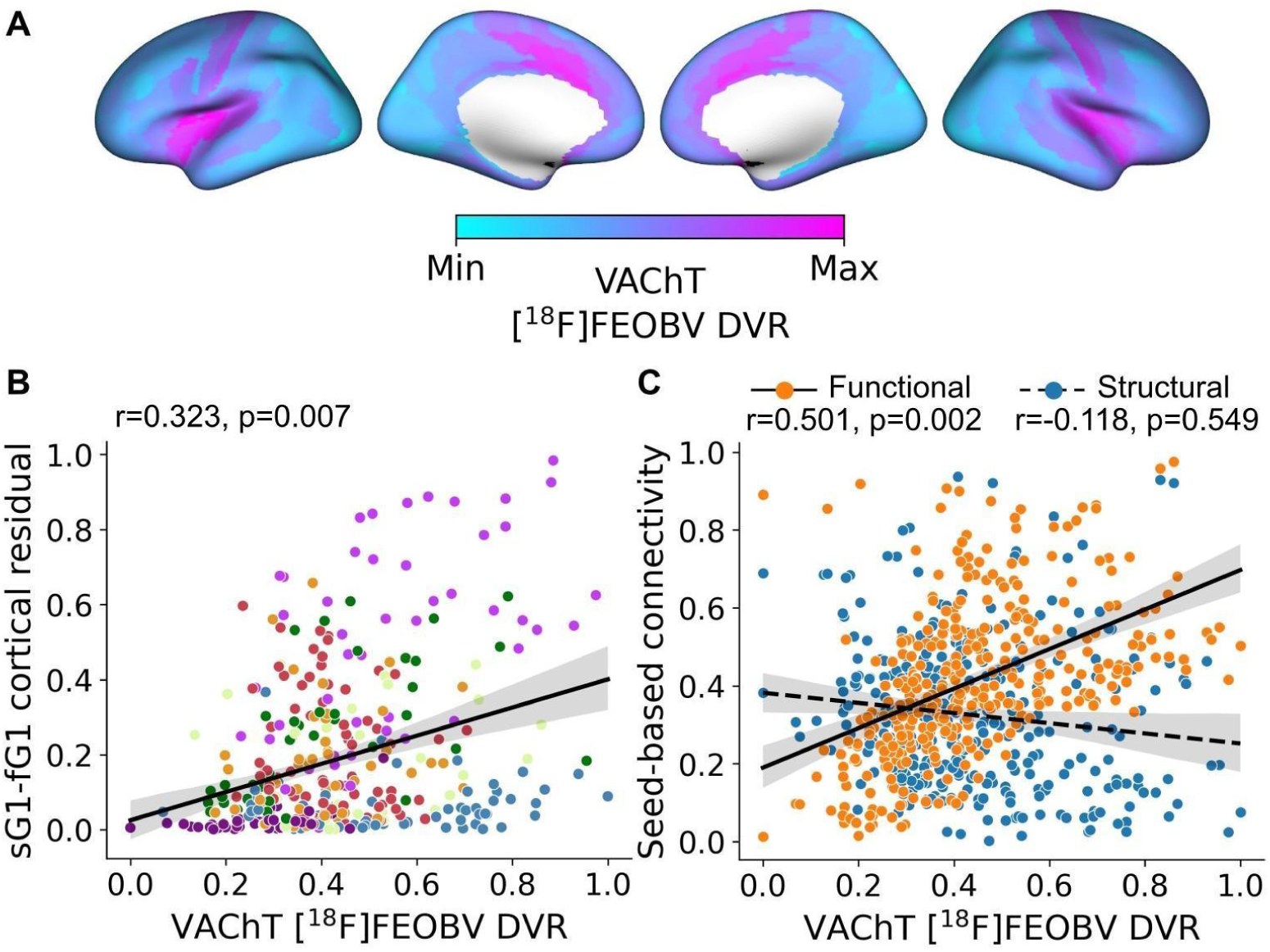
Multimodal gradients of basal forebrain (BF) connectivity in relation to *in vivo* molecular imaging of VAChT with [^18^F]FEOBV PET. (A) Average VAChT concentrations for 13 cognitively normal younger adults reveal the density of presynaptic cholinergic terminals across the cortical surface. (B) The spatial relationship of cortical VAChT concentrations (panel A) with the cortical map encoding BF structure-function tethering (Fig 2F). Each point in the scatter plot represents cortical parcels based on HCP-MMP 1.0 parcellation^29^ color coded according to the Yeo networks^25^ (Fig. 3B). (C) The spatial relationship of cortical VAChT concentrations (panel A) with seed-based connectivity for the BF structural (blue) and functional (orange) datasets.

Why might BF structural and functional connectivity diverge in cortical regions more densely innervated by BF cholinergic input? One possibility is that cortical areas expressing lower tethering between BF structural and functional connectivity tend to receive inputs from more highly branched cholinergic neurons. Under this ‘arborization gradient’ account, these cortical areas would exhibit strong BF functional connectivity and high VAChT concentration from multiple converging collateral arbors, but relatively weak structural connectivity due to greater diffusivity in the ascending tracts. To test this hypothesis, we computed seed-based connectivity of the BF with each HCP-MMP cortical parcel for both the rsfMRI (Pearson correlation r-values) and dMRI (streamline counts) datasets, and then projected these to the cortical surface. We then examined the spatial relationship of cortical VAChT concentration with the seed-based estimates of BF streamline counts and resting state correlations. Consistent with an arborization gradient, we observed a significant positive relationship of cortical VAChT concentrations with BF resting state correlations (R=0.501, p_spin_=0.002), but not streamline counts (R=-0.118, p_spin_=0.549; Fig. 4C). Moreover, the relationship of cortical VAChT with BF resting state correlations was significantly different than that observed for BF streamline counts (average difference between correlation coefficients = 0.608, 95% CI [0.727, 0.493], p_boot_<0.001, as revealed by bootstrap analysis), with greater divergence between BF resting state correlations and streamline counts in cortical parcels with higher VAChT concentrations.

We noted that the observed arborization gradient tends to differentiate cortical areas according to their distance from the BF nuclei, implying that arborization may reflect a biophysical constraint on cholinergic projections. The brain organizes its connections to optimize communication while minimizing the biological resources required for these connections, or ‘wiring cost’^51–53^. Neurons with long-range projections which extend from one brain region to another therefore often have fewer axonal branches than neurons with shorter-range projections. The BF cholinergic neurons may therefore balance longer distances traveled by their ascending axonal projections with fewer axonal arborizations. To test this hypothesis, we used the diffusion tractography data to compute the average white matter fiber length from the BF ROI to each cortical parcel (Methods). The fiber length estimates were then projected to the cortical surface (see Methods, Fig. 5A) and compared to BF structure-function tethering using spin tests^48,54^. Consistent with a distance/arborization tradeoff, we detected a significant negative relationship where cortical areas receiving longer fibers tend to exhibit a connectivity pattern of weak arborization, i.e. higher structure-function tethering (R=-0.280, p_spin_=0.027, Fig. 5B) and lower VAChT concentration (R=-0.483, p_spin_=0.004, Supplemental Fig. 6). In a distance/arborization tradeoff model, cortical areas at greater distances from the BF would require innervation from a larger number of individual BF neurons due to lower branching per neuron. This may necessitate greater numbers of BF streamlines to innervate distal cortical areas. Consistent with this prediction, we found that cortical areas innervated by longer fibers tend to receive higher BF streamline counts (Fig. 5C; R=0.455, p_spin_=0.0003), whereas fiber length was less related to the strength of BF resting state correlations (R=-0.226, p_spin_=0.455). Moreover, the relationship of fiber lengths with BF streamline counts was significantly different than that observed with BF resting state correlations (difference between correlation coefficients= -0.771, (95% CI [-0.664, -0.881], p_boot_<0.001). In sum, greater divergence between BF structural and functional connectivity, i.e. higher arborization, was observed in cortical parcels with higher VAChT concentration, stronger resting state correlations with BF, fewer direct streamlines, and shorter distances from BF.

**Fig. 5.**
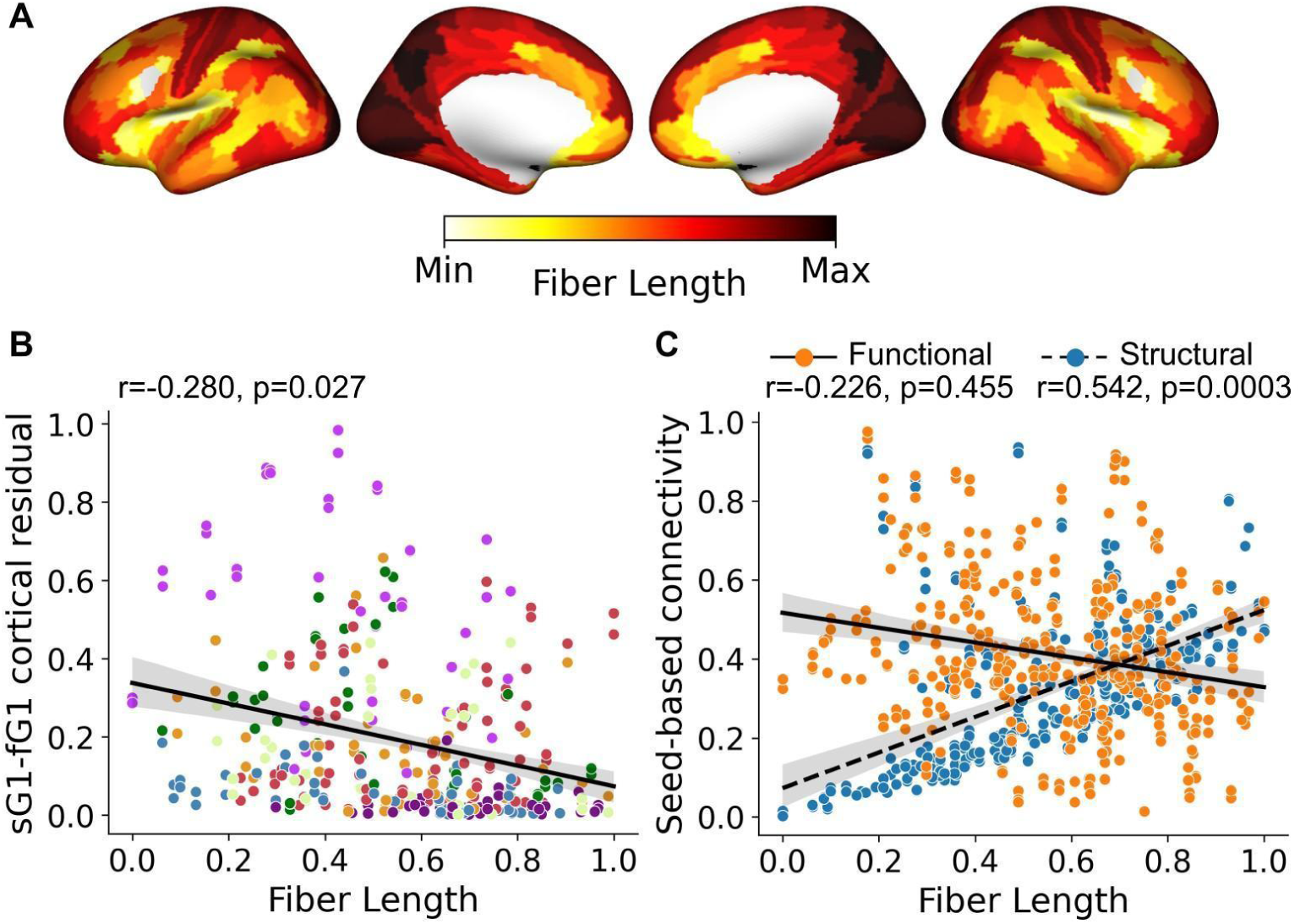
Multimodal gradients of basal forebrain (BF) connectivity in relation to diffusion tractography estimates of white matter fiber lengths. (A) Average white matter fiber lengths for 173 cognitively normal younger adults reveal cortical areas receiving the longest BF projections located in the primary visual and somatomotor cortices. (B) The spatial relationship of BF white matter fiber lengths (panel A) with the cortical map encoding BF structure-function tethering (Fig 2F). Each point in the scatter plot represents cortical parcels based on HCP-MMP 1.0 parcellation^29^ color coded according to the Yeo networks^25^ (Fig. 3B). (C) The spatial relationship of BF white matter fiber lengths (panel A) with seed-based connectivity for the BF structural (blue) and functional (orange) datasets.

If more individual BF neurons are needed to provide innervation over longer distances due to lower branching, this pattern would be consistent with higher wiring costs in these regions. We therefore next computed an estimate of BF wiring cost^22,23^ by multiplying the average fiber length with the total number of streamlines received from the BF (Fig. 6A, Supplemental Table 2) for each HCP-MMP cortical parcel. In this fiber distance-weighted structural connectivity map (Fig. 6B), the further away and the higher the streamline counts, the larger the wiring cost. Expectedly, the highest wiring costs were observed in cortical areas which receive input from cholinergic neurons predicted to have the lowest branching, i.e. primary visual and somatomotor cortex. These cortical areas also tend to have the farthest geodesic distances from the BF (Fig. 6C).

**Fig. 6.**
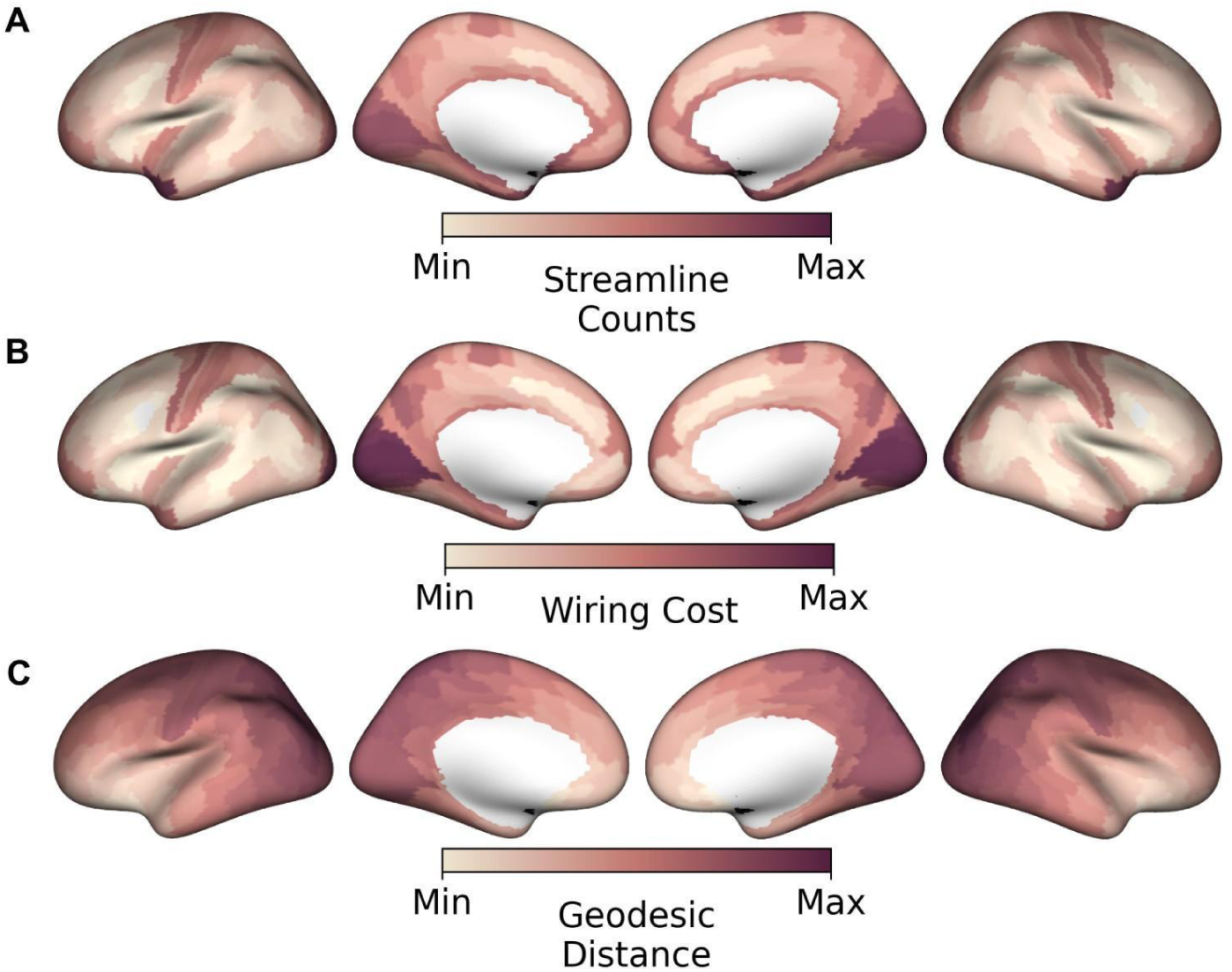
Cortical wiring costs and geodesic distances for basal forebrain (BF) projections. (A) Streamline counts representing the number of BF streamlines reaching each of the HCP-MMP cortical parcels^29^. (B) Wiring costs represent the BF white matter streamline counts for each cortical parcel weighted by their average fiber lengths^45^, (Fig 5A). Wiring costs are highest in primary visual and somatomotor cortices. (C) Geodesic distances from the BF to each parcel on the cortical surface.

### Cellular evidence for a gradient of arborization in BF cholinergic neurons

Thus far, the distance/arborization tradeoff observed in humans implies that the branch complexity of individual cholinergic neurons is shaped by a combination of the physical proximity and/or function of its cortical targets. However, a limitation of *in vivo* dMRI and rsfMRI techniques is that neither can resolve single cell axonal branching of cholinergic neurons. To test the arborization gradient hypothesis more directly, we leveraged a whole brain atlas of mouse BF cholinergic projections^2^ derived from cell-type specific anterograde viral labeling of individual BF cholinergic soma and axonal projections (Supplemental Table 1, Supplemental Fig. 7A). For each neuron, the targets of its axonal branches were labeled using the Allen Mouse Brain Atlas^55^, and classified according to whether any one of their branches targeted transmodal (iso)cortical areas. We identified 7/50 neurons fitting this criterion (Fig 7A). The Allen Mouse Brain Atlas^56^ annotations for the target areas include infralimbic and cingulate cortical areas within the mouse homologue of the salience network^57,58^. The remaining 43 neurons target unimodal sensory, motor or subcortical areas.

**Fig. 7.**
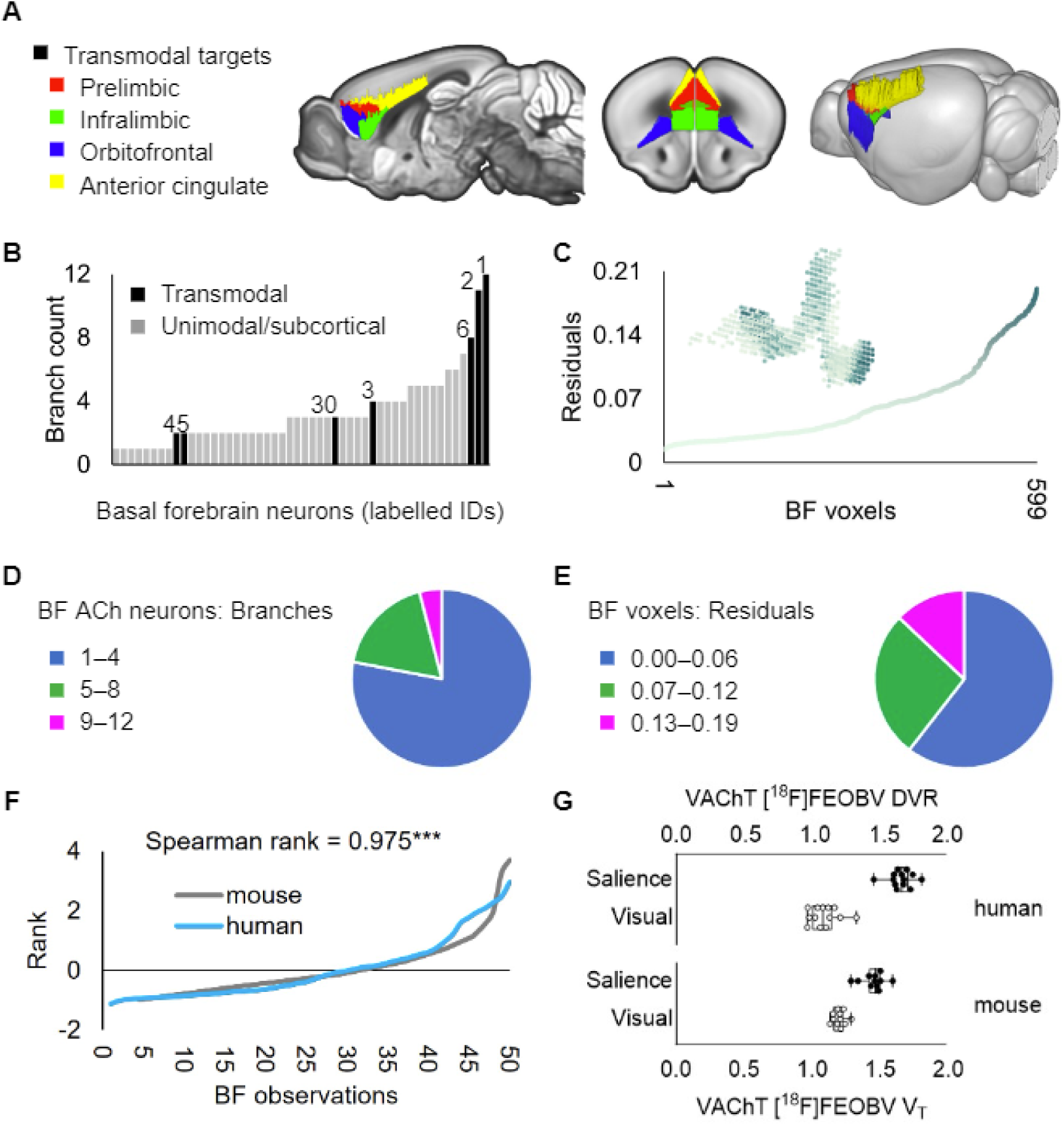
A cortical gradient of basal forebrain (BF) arborization in mouse and human. (A) In a whole brain atlas of mouse BF cholinergic neurons^2^ (n=50), 7/50 labeled neurons targeted transmodal (iso)cortical areas (color coded on sagittal, coronal and 3D renders of the Allen Mouse Brain Atlas^47^ common coordinate framework v3. (B) The number of distinct brain-wide targets for each of the 50 BF cholinergic neurons (branch counts) were ranked low to high. Black bars denote neurons targeting transmodal areas (panel A) and numerical label IDs correspond to individual neurons in supplemental Fig 7 of Li et al.^47^ . (C) The residual values encoding BF structure-function tethering in humans across 599 voxels of the BF region-of-interest (inset) were ranked low to high. (D and E) For mouse and human, distributions of neuronal branching and voxel residuals were split into tertiles to examine the concentrations of values (higher versus lower) relative to the total number of observations. (F) The rank ordered residuals across 599 BF voxels (downsampled to 50, blue) are superimposed over the branch counts of 50 mouse neurons (light gray) shown in panel B. (G) Comparisons of *in vivo* [^18^F]FEOBV PET data between ventral attention and visual networks (human) and between salience network regions (panel A) and a visual cortical control ROI (wild type mice, *n*=11).

We next counted the number of distinct brain-wide targets for each of the 50 BF cholinergic neurons. There was substantial variation in branch counts, ranging from neurons with only a single target to neurons with up to 12 different targets across the brain (Fig 7B). A permutation test comparing the number of branches between these two sets of neurons (transmodal targeting versus unimodal/subcortical targeting) revealed significantly higher branch counts in the transmodal targeting subset (10,000 permutations, t=3.2, p_perm_=0.004). To compare the distributions of BF neuronal branching in mice with our metric of BF neuronal branching in humans (structure-function tethering), we sorted the BF residuals (Figure 7C inset) from low to high across the 599 voxels comprising the full BF ROI. Consistent with mice, the distribution was right-skewed with a small concentration of higher residual values and a long left tail of smaller values (Fig 7C). For mouse and human, we split the respective distributions of neuronal branching and voxel residuals into tertiles to examine the concentrations of values (higher versus lower) relative to the total number of observations. For mice, the highest tertile contained only a small percentage (4%) of neurons with > 8 branches (Fig. 7D). Similarly for humans, the highest tertile contained a relatively small percentage (13%) of BF voxels (Fig. 7E). In both species, BF cholinergic neurons with very high branching thus represent a relatively smaller proportion of the total neuronal population. The shapes of the continuous distributions for single cell branch counts (Fig 7B) and for voxel residuals (Fig 7C) were also highly similar to one another (Spearman’s rank correlation = 0.975; Fig 7F), despite the fact that they originate from separate species and measurement techniques.

To bridge the gap between measures of central cholinergic innervation in human and mouse, we acquired *in vivo* [^18^F]FEOBV PET data in control mice (*n*=11, Supplemental Table 1) and quantified VAChT concentrations in the salience network regions (Fig. 7A) identified by the prior anterograde viral tracing study^2^ and a control region of interest comprising Allen Mouse Brain Atlas annotations for the visual cortex (see Methods, Supplemental Fig. 7B). In humans, we extracted [^18^F]FEOBV PET measures of VAChT from the homologous ventral attention and visual networks, as defined from the a priori Yeo^35^ networks (Fig. 3B). In both mice and humans, we found that VAChT concentrations were significantly higher in salience network hubs compared to the visual cortex (Fig. 7G; mouse: t_10_=11.54, p<0.0001; human: t_12_=17.19, p<0.0001). These findings provide further translational evidence that BF cholinergic neurons exhibit an arborization gradient which is shaped by the function and physical distance of their cortical targets.

To emphasize the features capturing the BF cholinergic arborization gradient in humans, we generated a cortical surface integrating measures of (1) BF structure-function tethering, (2) cortical VAChT concentration, and (3) fiber lengths of BF white matter projections (Methods, Fig. 8A). The highest convergence of these three features selectively colocalizes midcingulo-insular hubs of the ventral attention network (white boundaries). Altogether, our findings accord with a gradient of BF cortical cholinergic innervation which captures both modular (more neurons, fewer branches) and diffuse (fewer neurons, more branches) properties of functional integration^5^, depending on the subregional point of origin within the BF and the cortical targets (Fig. 8B).

**Fig. 8.**
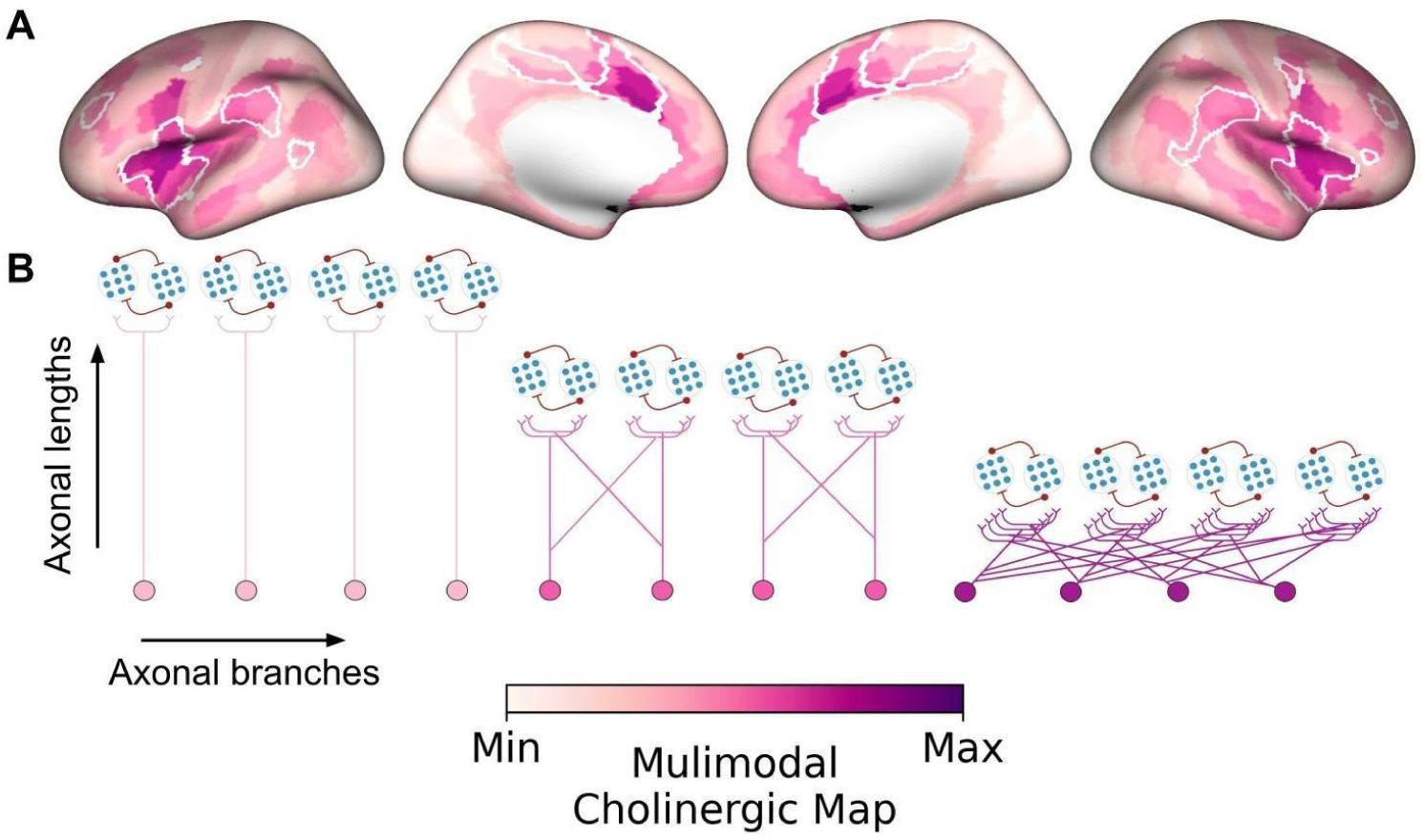
Multimodal map of the human basal forebrain (BF) cholinergic innervation. (A) A cortical surface emphasizing the arborization gradient of BF cholinergic neurons was constructed by summing the intensity normalized maps encoding (1) BF structure-function tethering, (2) cortical VAChT concentration, and (3) fiber lengths of BF white matter projections. The tethering and fiber length maps were sign flipped such that maximum values on the scale reflect higher VAChT, lower tethering, and shorter fiber lengths. The highest convergence of these three features selectively colocalizes midcingulo-insular hubs of the ventral attention network (white boundaries derived from Yeo et al^25^). (B) Depending on the subregional point of origin within the BF (anteromedial or posterolateral) and cortical target, BF cholinergic neurons may exhibit either a modular (more neurons, fewer branches) or diffuse (fewer neurons, more branches) profile of arborization.

## Discussion

We examined how the structural organization of cholinergic BF projections relates to their functional integration in the human and mouse cortex. Our findings, derived from multimodal in vivo brain imaging and cell type specific tracing of BF cholinergic axons, imply that the branch complexity of cholinergic projections may reflect properties of the cortical areas they target. Moving up the cortical hierarchy, neuronal populations with both increasingly diverse repertoires of cortical-cortical connectivity and shorter projection distances from the BF receive input from increasingly branched BF cholinergic neurons.

We considered the spatial relationship of the BF arborization gradient (defined from the residuals between BF structural and functional connectivity) to an established set of the brain’s core resting state networks^35^. Our findings indicate a pattern of cortical cholinergic innervation where shifts in axonal branching align spatially with boundaries of cortico-cortical network hubs. Cholinergic neurons with the highest arborizations target hubs of the midcingulo-insular network. Moving down this arborization gradient, frontoparietal and temporal network hubs exhibit an innervation profile of intermediate branching. Finally, cholinergic neurons with the fewest arbors per cell appear to preferentially target unimodal sensory cortices. In this hierarchically organized projection pattern, cholinergic neurons with fewer branches would need to be more numerous than those with many branches in order to provide coverage of an equivalent cortical surface area. Consistent with this prediction, cortical areas which receive higher numbers of BF streamlines tend to be farther away from the BF in terms of fiber lengths (Fig. 5C). Moreover, when we split estimates of axonal branch counts into tertiles in both mouse (cells) and human (voxels), the lowest tertile comprised >60% of observations (Fig. 7D,E), while the uppermost tertile comprised <15%. Hence, the BF appears to be populated by a significantly larger proportion of neurons with relatively fewer branches.

What is the functional significance of a cholinergic arborization gradient in this hierarchical cortical architecture? Neuroimaging work examining the functional and structural connectivity of the brain’s cortico-cortical networks has consistently identified the midcingulo-insular in high level selection and enhancement of both exogenous (sensory) and endogenous sources of input^28–36^. These two streams of information are thought to reflect a fundamental division of labor in the brain’s network organization. Support for this division comes from work demonstrating that exogenous and endogenous networks tend to be functionally anticorrelated with one another^59–63^, reflecting their competitive interactions for representational dominance of the outside sensory world versus internal mental content. The midcingulo-insular areas are unique in that they exhibit positive temporal correlations with networks in both the exogenous and endogenous processing streams, implying a superordinate role in selecting between the two^36,64–66^. We also recently provided meta-analytic evidence that task-related midcingulo-insular activations, as measured by fMRI, exhibit among the strongest patterns of pharmaco-modulation by cholinergic agonists and task-related co-activations with the BF^67^. The profile of BF cholinergic arborization and strength of its modulatory regulation on midcingulo-insular cortex provide further evidence of its privileged supervisory status.

Cortical cholinergic signaling is thought to play a central role in promoting the selection and processing of relevant stimuli while suppressing competing irrelevant information^5,68–70^. Parallel lines of research in mouse and non-human primates has started to uncover the cortical circuit and computation which enable this core property of brain function^71^. The BF cholinergic inputs form an integral component of a canonical disinhibitory microcircuit expressed throughout the cortex, composed of pyramidal neurons and multiple classes of interneurons^72–80^, possibly organized at the columnar scale^81,82^. Targeted release of acetylcholine in this microcircuit may bias mutually suppressive responses of neuronal populations competing for attentional resources^71,83–85^, allowing one population to dominate over others. Moving up the cortical hierarchy, individual cholinergic neurons may require larger arborizations to bias competitive responses across increasingly large and spatially distributed populations of cortical neurons. We speculate that coalitions of BF cholinergic neurons may coordinate the release of acetylcholine^86,87^ along this arborization gradient. These transient coordinated patterns of signaling may act to flexibly segregate^88^ assemblies of neurons^89–92^ across the cortical hierarchy to form coherent attentional episodes^93^.

A diversity of branch complexity in BF cholinergic neurons may also account for differences in their vulnerability to aging and disease. Cell type specific labeling and transcriptomic analyses examining morphological and functional properties which increase a neuron’s vulnerability to age-related neurodegenerative disease such as AD have consistently demonstrated large axonal projections as a key risk factor^4,94–96^. The observed decreased structure-function tethering in the postero-lateral BF, and in particular the NbM, is consistent with neurons exhibiting large arborizations. This morphofunctional property of NbM cholinergic neurons may increase their vulnerability to dysfunction in the aging brain. In parallel, our observation that ventral attention network may receive input from the most highly branched NbM cholinergic neurons implies that these cortical areas might exhibit higher vulnerability to dysfunctional cholinergic signaling in the aging brain^97^. It is also notable that the BF cholinergic neurons with fewer arborizations, which our findings suggest primarily target the primary and somatosensory cortices, constitute projection zones which are relatively spared by pathology in early stages of AD^98^.

Our findings are subject to several important methodological considerations. First, the basal forebrain is a small subcortical structure with poorly defined anatomical boundaries. We therefore used a probabilistic atlas to localize its constituent nuclei. However, when using probabilistic BF atlases in combination with data collected at spatial resolutions typical of 3T structural (1.5 mm^3^) and functional MRI (3 mm^3^), aliasing of adjacent structures has been shown to systematically overestimate the BF gray matter^99^. To mitigate this issue, we used optimized MRI protocols to acquire high spatial resolution dMRI (1.05 mm^3^) and rsfMRI (1.6 mm^3^) at 7T ^100,101^. Second, our measures of structural connectivity were estimated using streamline tractography on diffusion-weighted imaging, which can be susceptible to false positives and negatives in certain brain areas^99^. It is therefore possible that the regional variation in BF structure–function correspondence is partly explained by regional variation in tractography performance. Another concern is the susceptibility-related spatial distortions near the BF region for fast readout scans, such as those used for the rsfMRI acquisitions by the HCP. Although corrected for using a separately acquired field map, these spatial distortions might lead to suboptimal probing of BF voxels in such data with possible contamination from white matter (WM) tissue and cerebro-spinal fluids (CSF). To limit the impact of the latter on the functional time series analysis, additional denoising using the average WM and CSF timeseries was performed. Finally, the atlas provided by Li et al.^2^ consists of cholinergic neurons sampled exclusively from anteromedial nuclei of the basal forebrain. Complete morphologies of individual basal forebrain cholinergic neurons are needed over larger samples of neurons, with greater coverage of the mouse basal forebrain nuclei, and with quantification of both axonal and dendritic arborizations^3,4,6^.

In sum, we demonstrate that multimodal gradients of BF connectivity reveal spatially inhomogeneous patterns of structure-function tethering in the cortex, with the lowest tethering in mid-cingulate and anterior insular cortical areas involved in salience detection and allocation of attentional resources throughout the brain. These cortical areas tend to be located proximal to the BF and receive disproportionately higher concentrations of cholinergic innervation. Our findings conform to a model in which this multimodal BF gradient is shaped in part by axonal branch complexity.

## Materials and Methods

We used high-resolution minimally pre-processed 7T MRI HCP data (n=173)^39^ and the existing stereotactic atlas of the BF ^40^ to build structural and functional connectomes. Any further pre- and postprocessing was done on the high-performance computing cluster of the Digital Research Alliance of Canada. Workflows were built using Snakemake^102^ with the full workflow available on GitHub (see data and code availability for specifics). Individual connectomes were averaged and reduced to a 2-dimensional *M*-by-*N* matrix describing the pairwise connectivity strength between *M* BF ROI voxels and *N* cortical regions^41^. The BrainSpace toolbox^44^ was used to capture the gradients which, as well as any further analysis, was done using Jupyter Notebooks^103^.

### Human Data Acquisition

High-resolution 7T dMRI and rsfMRI data were downloaded from the HCP data repository^24^. We used the minimally pre-processed data (described in ref 39^39^) consisting of 173 healthy subjects (69 male, 104 female) aged 22 to 35 years. The dMRI images were collected with a 1.05 mm^3^ isotropic voxel size, TR=7000 ms, TE=71.2 ms, b-values=1000, 2000 s/mm^2^, FOV=210 x 210 mm^2^. Resting-state fMRI images were collected with a 1.6 mm^3^ isotropic voxel size, TR=1000 ms, TE=22.2 ms, FOV=208 mm^2^, spanning 4 runs of 16-minute duration each, per subject. For anatomical imaging, two T_1_-weighted (T_1_w) scans were obtained using a three-dimensional (3D) magnetization-prepared rapid gradient-echo (MPRAGE)^104^ sequence with identical geometries and a 0.7 mm^3^ isotropic voxel size. Full details of the acquisition parameters can be found in the HCP S1200 release reference manual^105^.

### Basal Forebrain Mask

The BF ROI was created using the existing stereotactic atlas of the BF^40^. This stereotactic BF atlas is based on histological sections obtained from 10 postmortem brains. The magnocellular cell groups were delineated in each slice, 3D reconstructed and warped into the MNI single-subject reference space^106^. The atlas consists of 4 subregions of the BF defined in the nomenclature: Ch1-2, Ch3, Ch4, and Ch4p^107^. For each subregion, a stereotactic probabilistic map has a range of 0 to 10 indicating the number of brains containing the specific magnocellular cell groups in the given voxel. Note that this atlas is not left, right symmetrical; hence, our BF ROI is asymmetrical. Our BF ROI is created by thresholding these subregion masks to 50% first and then combining all to get a mask covering full BF. This BF ROI mask was then warped into MNI152 non-linear 6^th^ generation atlas (MNI152Nlin6Asym)^106^.

### Structural Connectivity Reconstruction

Diffusion tractography was performed to get a connectivity matrix for diffusion data. As part of the minimal preprocessing pipeline data release, all subjects underwent FreeSurfer processing (v5.3.0-HCP)^108^. The BF ROI mask was then first resampled and transformed to the individual subjects’ minimally preprocessed volume space (0.7mm^3^). Volumetric cortical labels were built by mapping the HCP-MMP 1.0 surface parcellation^41^ using Connectome Workbench’s ribbon-constrained *label-to-volume-mapping* function and FreeSurfer-derived surfaces. The BF ROI voxels were used as seeds, and the 180 cortical regions in each hemisphere were combined and used as targets to perform probabilistic tractography using FSL’s *probtrackx*^109^ with 5000 streamlines per BF ROI voxel. The resulting probability maps (Supplemental Fig. 1C) in the BF quantified the number of streamlines that reached each target. The maps were resampled to MNI space^110^ and 1.6 mm^3^ resolution to match the functional connectivity data and reduced to a 2-dimensional *M*-by-*N* matrix, where *M* represents the voxels in the BF ROI (599 voxels) and *N* is the cortical targets (180 each hemisphere) with their corresponding number of streamlines. Additionally, the ‘wiring cost’ connecting the BF ROI with the cortex was calculated by multiplying the number of streamlines with their average length for each cortical target (Gollo et al. 2018). The *M*-by-*N* connectivity feature matrix, as well as cortical wiring cost map for all 173 subjects were averaged to calculate the gradients and wiring cost, respectively.

### Functional Connectivity Reconstruction

First, the BF ROI mask in the minimally preprocessed volume space was resampled to the 1.6 mm^3^ isotropic voxel size of the rsfMRI data and added to the subject’s subcortical parcellation. A functional connectivity matrix was then created for each subject by calculating the temporal correlation between BF voxels and cortical ROIs. All four runs (i.e., two sets of 16 min. runs with posterior-to-anterior and anterior-to-posterior phase-encoding) of the minimally preprocessed and ICA-FIX denoised rsfMRI data^111^ were used. Since the BF ROI is not included in the dense timeseries provided by HCP, these were regenerated using the updated subcortical parcellation to include the BF ROI voxels for further processing. Subsequent processing included ROI-constrained subcortical smoothing to match the cortical sampling density using the scripts provided by HCP^108^, as well as additional signal filtering (i) based on the average WM and CSF timeseries using *ciftify*^112^ and (ii) by applying a Wishart filter as proposed previously^113,114^ to selectively smooth unstructured noise more than the structured blood oxygenation level-dependent signal. Average cortical ROI timeseries (concatenated across runs) were then extracted using the HCP-MMP 1.0 surface parcellation^41^. The 2-dimensional *M*-by-*N* functional connectivity matrices were constructed by calculating the Pearson’s correlation coefficient for each BF ROI voxel (*M*) to each of the cortical parcels (*N,* 180 each hemisphere) and averaged over all subjects to calculate group-wise gradients.

### Gradient Calculation

Connectivity gradients were calculated using the BrainSpace toolbox^44^. Group averaged connectivity matrices were used as input to the *GradientMaps* function, using the normalized angle kernel and diffusion map embedding approach. This nonlinear dimension reduction method transforms the connectivity matrix into a low-dimensional representation to construct connectivity gradients and their corresponding lambda values (i.e., explained variance)^25^. BF voxels that are characterized by similar connectivity patterns will have a gradient value closer together, whereas voxels with little or no similarity are farther apart. These gradients were then mapped back onto the BF voxel space to visualize continuous transitions in functional and structural connectivity patterns (Supplemental Fig. 2A).

In addition, gradient-weighted cortical maps were created by multiplying each row of the BF-cortical connectivity matrix with the corresponding gradient value of that BF voxel ^115^ (Supplemental Fig. 2B). The distribution of cortical gradient-weighted values was then decomposed into seven functional networks^35^ using the HCP-MMP 1.0 parcellation-based 7 Yeo networks as defined in ref 35 and 116^35,116^. These include the visual, somatomotor, dorsal attention, ventral attention, limbic, frontoparietal, and default mode network.

Residual values (structure-function tethering) for each of the BF voxels and HCP-MMP parcels were calculated based on the regression between the structural against the functional gradients for the BF and gradient-weighted cortical maps, respectively.

### Human [^18^F]FEOBV PET

[^18^F]FEOBV PET acquisition and preprocessing is described in Kanel et al.^38^ Each individual [^18^F]FEOBV PET image was intensity normalized to the subject’s supratentorial white matter uptake to create a parametric DVR [^18^F]FEOBV PET image^38^. The original PET atlases were transformed to *10k_fsavg* surface-space and parcellated to HCP-MMP 1.0^41^. The values of each cortical parcel encoding the relative concentration of cholinergic nerve terminals were rescaled^117^ and visualized on an inflated surface (Fig. 4A). Additional [^18^F]FEOBV maps^49,50^ were obtained from the Neuromap toolbox^54^ to examine the reproducibility of our results.

### Mouse [^18^F]FEOBV PET

All small-animal imaging procedures were conducted in accordance with the Canadian Council of Animal Care’s current policies and were approved by the University of Western Ontario’s Animal Care Committee (Animal Use Protocols: 2020-162 and 2020-163) and the Lawson Health Research Institute’s Health and Safety board.

High-resolution 9.4T MRI and [^18^F]FEOBV PET data were acquired at the Centre of Functional and Metabolic Mapping (Agilent Animal MRI Scanner, Bruker) and the Lawson Health Research Institute’s Preclinical Imaging Facility (Inveon DPET, Siemens Healthineers), respectively. Data were acquired in 11 (5 male, 6 female) control mice, either C57BL/6J or VAChT^flox/flox118^ at 6 months of age.

*In vivo* dynamic [^18^F]FEOBV PET data were acquired as 150-minute list-mode emission scans, performed with a timing window of 3.432 ns and a 350–640 keV discrimination energy range. Mice were anesthetized (1.5-2.0% isoflurane in 1.0 L/min O_2(g)_) and then injected intravenously with a ∼20 Megabecquerel (MBq) dose of [^18^F]FEOBV. Data were binned into consecutive frames of increasing duration (*n*_frames_ = 43), consisting of 15 frames of 40 seconds, followed by 28 frames of 300 seconds. An OSEM3D algorithm with two iterations and 18 subsets was used to reconstruct [^18^F]FEOBV PET images with the following parameters: matrix dimensions: 128 mm × 128 mm × 159 mm; voxel resolution: 0.776 mm × 0.776 mm × 0.796 mm.

Given that (a) VAChT is expressed throughout cerebral^2^, subcortical, and cerebellar grey matter regions of the mouse brain^119^, and (b) the spatial resolution of microPET cannot resolve [^18^F]FEOBV uptake within small white matter structures, no anatomical reference region was used for intensity normalization of mouse [^18^F]FEOBV PET data. Rather, parametric volume of distribution (V_T_) images were estimated for each mouse by fitting the whole-brain [^18^F]FEOBV time-activity curves with the multi-time point graphical analysis methodology described by ^120^, where an image-derived input function (IDIF) served as the reference. The IDIF was obtained from a volume of interest placed in the lumen of the left ventricle of the mouse heart^121^, and the start of the linear portion (t*) was determined from the curve fitting, where t* = ∼11.33 minutes.

Pre-processing of mouse [^18^F]FEOBV V_T_ images was performed using SPM12 (https://www.fil.ion.ucl.ac.uk/spm/software/spm12/) in MATLAB R2020b. Preparation of [^18^F]FEOBV V_T_ images included resizing, centering, and re-orienting the images to follow the right-hand coordinate system (RAS). Each V_T_ image was then coregistered to its corresponding native space MRI image using normalized mutual information. Finally, all [^18^F]FEOBV V_T_ images were cropped to the native space MRI bounding box dimensions and nonlinearly warped to the study-specific anatomical mouse brain template (see below). To parcellate mouse [^18^F]FEOBV V_T_ images, we performed non-linear registration of the study-specific mouse brain template to the Allen Mouse Brain Atlas^55^.

For anatomical MRI, mice were anesthetized (1.5-2.0% isoflurane in 1.5 L/min O_2 (g)_) and then positioned in the center of a 30 mm RF volume coil (Bruker). Four iterations of magnetization transfer (MT)-weighted spoiled gradient-recalled echo (GRE) images were acquired using a fast low angle shot 3D pulse sequence with the following parameters: FOV = 16 mm x 16 mm x 10 mm; matrix dimensions: 128 x 128 x 8; voxel resolution = 0.125 mm^3^, slice thickness = 10 μm; TR = 30 ms; TE = 2.77 ms; FA = 9°; off-resonance saturation = 4.5 kHz (Gaussian pulse = 10 ms duration); effective FA = 900°^122^.

Pre-processing of mouse anatomical brain images was performed using SPM12 (https://www.fil.ion.ucl.ac.uk/spm/software/spm12/) in MATLAB R2020b. For each mouse, the anatomical brain images (*n*=4) were first averaged, then resized, centered, and re-oriented to follow the RAS. Using normalized-cross correlation, the images were then coregistered to the mouse brain template from Hikishima et al.^123^, bias corrected using MICO^124^, and then segmented using tissue prior probability maps for gray matter, WM, and CSF tissue compartments. For each mouse, the tissue segments from this step were combined to mask out the surrounding skull and tissue. Finally, the gray and white matter segmentations were used to create nonlinear transformations for diffeomorphic registration to a population specific average template using geodesic shooting ^125^.

### Geodesic Distance

Geodesic distance along the cortical surface was calculated using the geodesic library (https://github.com/the-virtual-brain/tvb-gdist) based on the algorithm that approximates the exact distance along the shortest path between two nodes (or vertices) on a triangulated surface mesh^126^. An average BF seed node was created for the left and right hemispheres separately by (i) projecting the BF mask onto the *59k_fs_LR* white matter surface of the individual subjects using Connectome Workbench’s *volume-to-surface-mapping* function, (ii) averaging across all subjects to get a probability map, (iii) resampling to the *10k_fsavg* surface-space as suggested by the HCP study (https://wiki.humanconnectome.org) and (iv) then by thresholding at 0.5 to obtain a final binary BF seed on the cortical surface. A distance value was then assigned to each cortical vertex based on the minimum geodesic distance along the *10k_fsavg* pial surface to the BF seed node to avoid the medial wall. To match with the resolution of the cortical connectivity results, the geodesic distance map was parcellated using the HCP-MMP 1.0 atlas as implemented in the neuromaps toolbox^54^, and rescaled to values between 0 and 1^117^.

### Statistical analyses

Permutation tests with surrogate maps^127^ were used to compute statistical significance for the BF gradients and the distribution of residuals. BF gradient values for structural and functional connectivities were first rescaled between 0 and 1 and Euclidean distance was used to calculate the distance matrix among all voxels within the original BF ROI^117^. Variograms were then permuted (N=1000) using the *SurrogateMaps* function implemented in the BrainSpace toolbox^44^. Parameters were adjusted in the case of a suboptimal fit compared to the empirical data (pv=60, random_state=1234). The final variograms were used to build and compare null probability for the mean and CoV between BF subregions.

To assess the stability of structural and functional connectivity gradients and the gradient of their tethering, we performed split-half and leave-one-out cross-validation analyses across the 173 individuals in the sample (Supplemental Fig. 3). For each split, or subject left out, the model was trained on the remaining individuals, and the predicted gradient was compared to the observed gradient for the left-out individual.

Spin tests^48^, as implemented in the neuromaps toolbox^54^, were used to compare cortical maps based on N=10k permuted maps. All cortical maps were parcellated using the HCP-MMP 1.0 atlas^44^ and values were rescaled between 0 and 1^117^. In addition to the spin tests, a bootstrapping analysis was performed to quantify the difference between structural and functional connectivity and their correlation with [^18^F]FEOBV PET maps and fiber length. Here, bootstrapping was applied 10k times (by randomly selecting sets of regions during each iteration) to build a null probability of correlation coefficients for statistical inference based on the observed difference between the two connectomes.

### Data and code Availability

For origin and use of the multiple datasets described in this paper, see Supplemental Table 1. The Human Connectome (HCP) project dataset is available at http://www.humanconnectomeproject.org/. The human [^18^F] FEOBV PET data was derived from Kanel et al.^38^ and neuromaps^37^. The mouse [^18^F] FEOBV PET data was collected by K.M.O. and is available upon request. The viral tracing data were derived from Li et al.^2^ and are available in their supplemental materials (SI Appendix Fig 7). The workflow for reconstructing structural connectivity matrix from the HCP data and a subcortical region of interest is available at https://github.com/sudesnac/diffparc-smk ^128^; and the functional connectivity workflow at https://github.com/khanlab/subcorticalparc-smk ^129^. All other code used to conduct the reported analyses and create the figures are available at https://github.com/sudesnac/HumanBF-Connectivity ^130^.

## Author Contributions

Author contributions: S.C., A.R.K. and T.W.S designed research; S.C., R.A.M.H., K.MO., and A.R.K. performed research; A.R.K., R.A.M.H., and K.M.O. contributed new reagents/analytic tools; PK and K.M.O conducted preprocessing of PET images; PK also reviewed and critiqued paper; M.A.M.P. and V.P. provided the mouse lines; S.C. and R.A.M.H. analyzed data; and S.C. and T.W.S. wrote the paper

## Classification

Biological Sciences, Neuroscience

## Acknowledgments

RH was supported by a BrainsCAN postdoctoral fellowship as well as a Marie Skłodowska-Curie Actions Postdoctoral Fellowship (101061988). This research was enabled in part by the support provided by the Digital Research Alliance (https://alliancecan.ca), and the Natural Sciences and Engineering Research Council (TWS). Data was provided in part by the Human Connectome Project, WU-Minn Consortium (Principal Investigators: David Van Essen and Kamil Ugurbil; 1U54MH091657) funded by the 16 NIH Institutes and Centers that support the NIH Blueprint for Neuroscience Research; and by the McDonnell Center for Systems Neuroscience at Washington University. The Human PET data used in the analysis was supported by the National Institutes of Health [P01 NS015655, RO1 NS070856, P50 NS091856, P50 NS123067].

