## Supplemental Tables and Figures for "Multimodal gradients of basal forebrain connectivity across the neocortex"

### Supplemental Materials:

**Supplemental Table 1. Summary of the datasets.** Demographic details of all the datasets used in this study. Ages presented as mean (standard deviation), where applicable.

| Species | Reference | Imaging | Imaging materials | Use in Chakraborty et al. | Strain | Age (y or m) | Sex (M:F) |
|---|---|---|---|---|---|---|---|
| Human | HCP | 7 Tesla MRI | dMRI | gradient calculation, residual analysis | N/A | 22-35 y | 69:104 |
| | | | rs-fMRI | | | | |
| Human | Kanel | PET | [$^{18}$F]FEOBV | principal correlation analysis of structure-function tethering and cholinergic innervation | N/A | 24.5(4.9) y | 10:3 |
| Human | Aghourian | PET | [$^{18}$F]FEOBV | replication correlation analysis of structure-function tethering and cholinergic innervation | N/A | 66.8(6.8) y | 5:13 |
| Human | Bedard | PET | [$^{18}$F]FEOBV | replication analysis of structure-function tethering and cholinergic innervation | N/A | 68.3(3.1) y | 4:1 |
| Human | Tuominen | PET | [$^{18}$F]FEOBV | replication analysis of structure-function tethering and cholinergic innervation | N/A | 37(10.2) y | 3:1 |
| Mouse | Li | SIM | AAV-CAG-flex-GFP | unimodal versus transmodal cholinergic neuron branch counts with retrospective mouse data | *ChAT*-ires-Cre | 3-6 m | N/A |
| Mouse | N/A | microPET | [$^{18}$F]FEOBV | cross validation of human cortical cholinergic innervation with prospective mouse data | *VAChT*$^{flox/flox}$ | 6 m | 3:3 |
| | | | | cross validation of human cortical cholinergic innervation with prospective mouse data | C57BL/6J | | 2:3 |

**Supplemental Table 2: Streamline counts.** The excel file detailing the streamlines received at each of the cortical parcels (left and right combined) from the BF (all voxels) available at https://github.com/sudesnac/HumanBF-Connectivity/blob/main/data/Diff_streamline-counts_summed_seed-BASF_voxels.xlsx

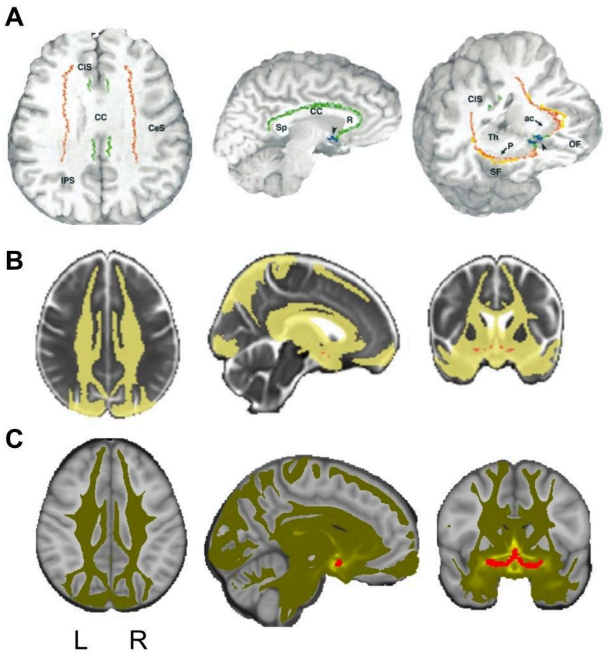


**Supplemental Fig. 1: Basal forebrain (BF) cholinergic white matter pathways.** (A) Retrograde and anterograde labeling of cholinergic (ChAT+) fibers from the nucleus basalis of Meynert in human postmortem data[1]. The basal forebrain nuclei are visible on the coronal oblique section. Two core cholinergic BF projections were identified: a medial cingulum pathway (green) and a lateral capsular pathway (red) with a perisylvian division (orange). (B) *In vivo* diffusion MRI tractography of the BF nucleus basalis of Meynert in a large sample of adults (N=262)[2]. Note that the medial and lateral tracts identified in vivo closely recapitulate the post-mortem tracing results. (C) Diffusion tractography results from the present dataset. The strongest weighting corresponds to medial (cingulum) and lateral (capsular) BF pathways.

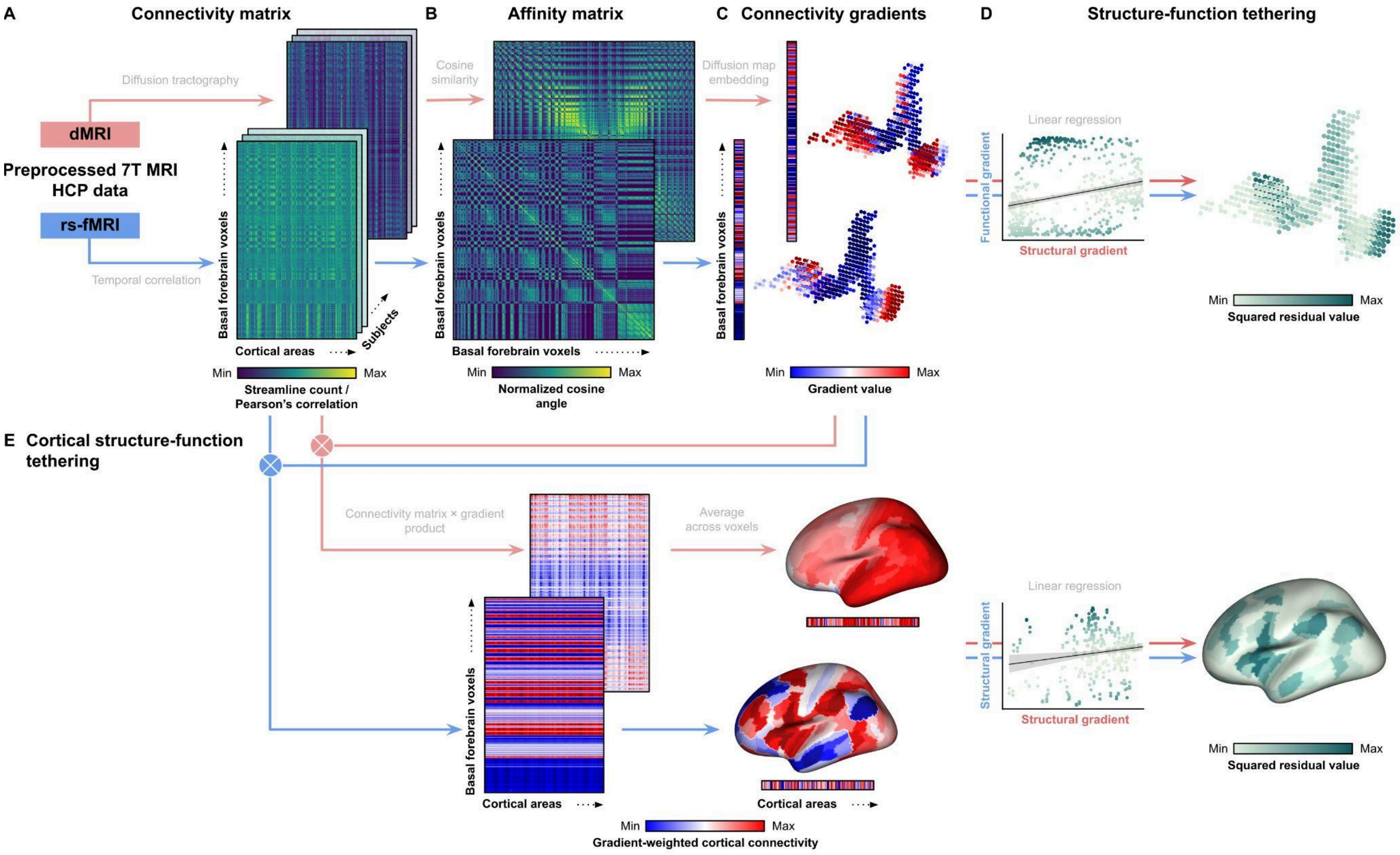

A
Connectivity matrix
B
Affinity matrix
C
Connectivity gradients
D
Structure-function tethering
Diffusion tractography
dMRI
Preprocessed 7T MRI
HCP data
rs-fMRI
Temporal correlation
Basal forebrain voxels
Cortical areas
Subjects
Min
Max
Streamline count /
Pearson's correlation
Cosine
similarity
Basal forebrain voxels
Min
Max
Normalized cosine
angle
Diffusion map
embedding
Min
Max
Gradient value
Linear regression
Functional gradient
Structural gradient
Min
Max
Squared residual value
E
Cortical structure-function
tethering
Connectivity matrix × gradient
product
Average
across voxels
Cortical areas
Cortical areas
Min
Max
Gradient-weighted cortical connectivity
Linear regression
Structural gradient
Structural gradient
Min
Max
Squared residual value

**Supplemental Fig. 2. Calculation of basal forebrain (BF) structure-function tethering.** (A) Diffusion MRI (dMRI; red arrows) and resting-state fMRI (rs-fMRI; blue arrows) data were processed separately to extract structural and functional connectivity matrices. dMRI data were used to reconstruct streamlines via diffusion tractography, while rs-fMRI data were analyzed for temporal correlations between BF voxels and cortical regions. (B-C) BF gradients were computed to demonstrate principal axes of connectivity variability among BF voxels, highlighting significant patterns based on inter-voxel similarity using the normalized cosine angle. (D) Linear regression was conducted between the first structural gradient (from streamline counts) and the first functional gradient (from Pearson's correlation strength) to derive voxel-wise residual values, measuring the degree of 'tethering'. (E) Gradient-weighted cortical maps were created by multiplying each row of the initial connectivity matrices with the corresponding principal gradient value, then averaging these rows to produce a single cortical representation of each gradient. Linear regression was then applied between the structural and functional gradient-weighted cortical maps to derive residual values for each cortical area.

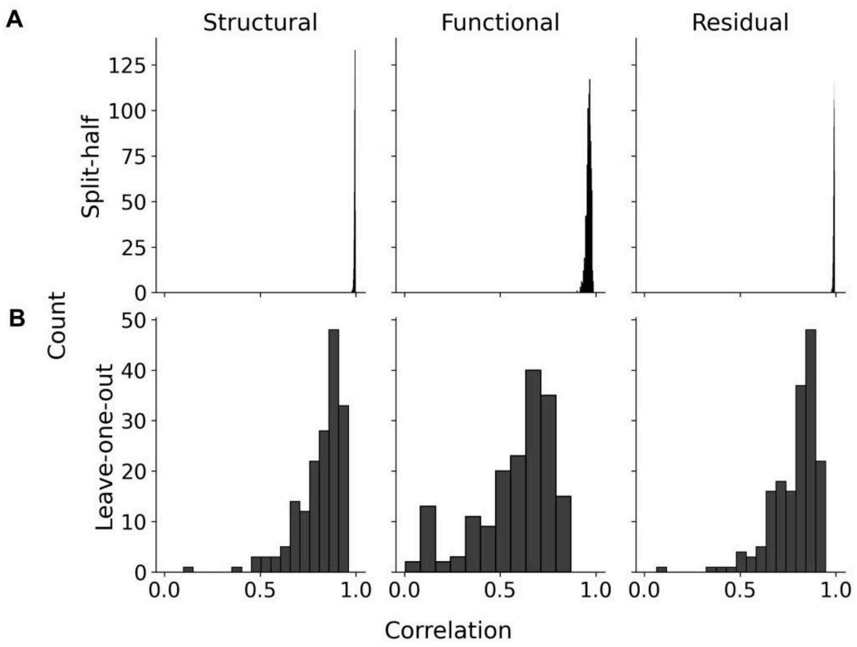


**Supplemental Fig. 3. Calculation of split-half and leave-one-out cross-validation of basal forebrain (BF) gradients.** (A) For each split (N=1000), the model was trained on the other half, and the predicted gradient was compared to the observed gradient for the left-out half. (B) For each subject left out (N=173), the model was trained on the remaining individuals, and the predicted gradient was compared to the observed gradient for the left-out individual.

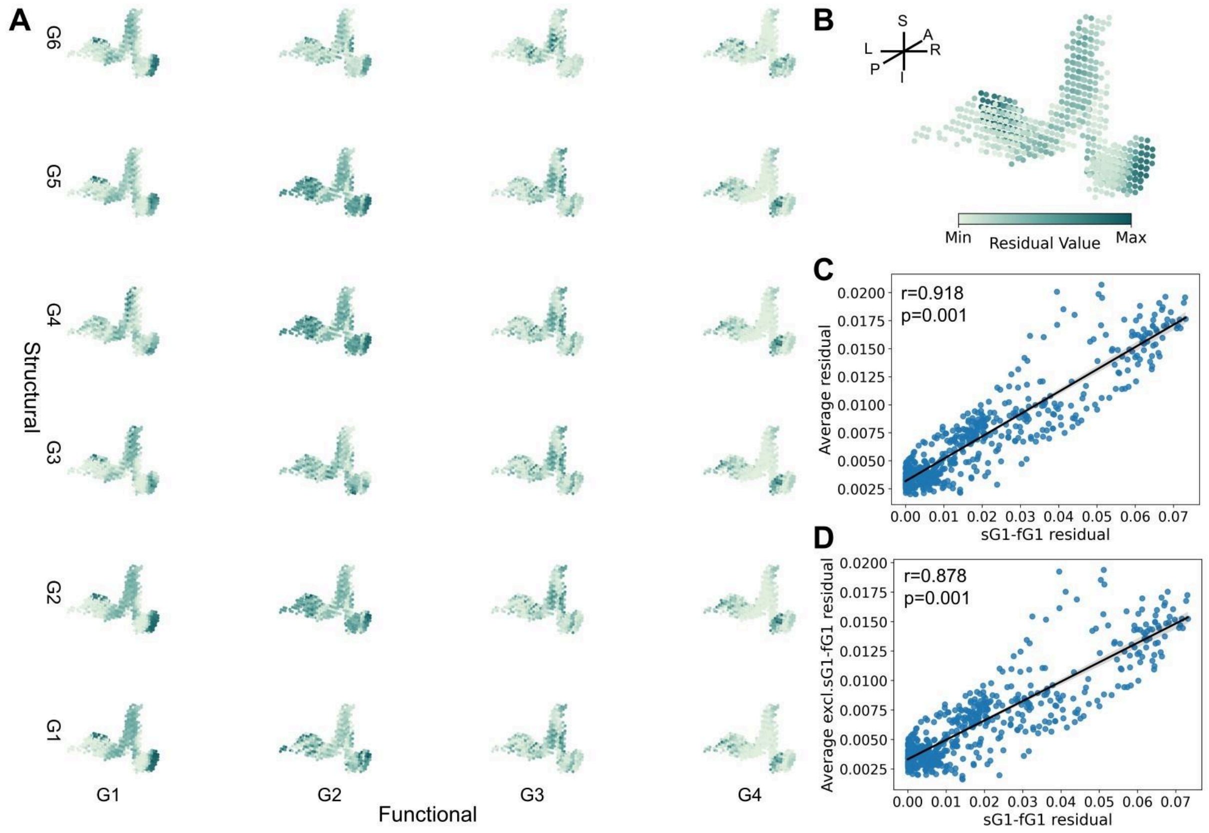


**Supplemental Fig. 4. Calculation of structure-function tethering for all gradients falling above the knee points for their corresponding structural and functional gradients.** (A) The knee-point[3] corresponded to sG6 for structural gradients and fG4 for functional gradients, thus forming 6 x 4 = 24 structure-function gradient pairs. (B) The average of the 24 residual maps qualitatively is highly similar to the sG1-fG1 residual map. (C) The Pearson between sG1-fG1 residual map (x-axis) and average of the 24 residual maps (y-axis). (D) The Pearson between sG1-fG1 residual map (x-axis) and average of the 23 residual maps, excluding sG1-fG1 residuals (y-axis).

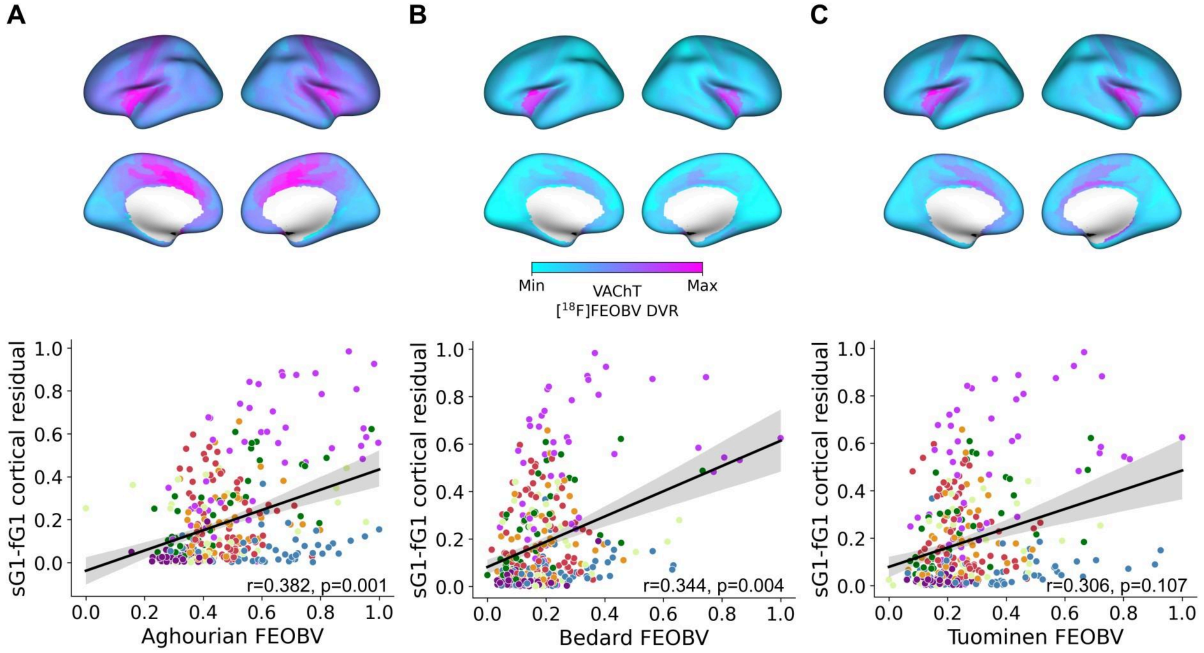


**Supplemental Fig 5. Relation of basal forebrain (BF) structure-function tethering to three publicly available [$^{18}$F]FEOBV PET maps.** (A) Aghourian et al[4] (B) Bedard et al.[5] (C) unpublished data provided to neuromaps by Tuominen et al.[6] Each point in the scatter plots represents cortical parcels based on HCP-MMP 1.0 parcellation color coded according to the Yeo networks[7] (Fig. 3B).

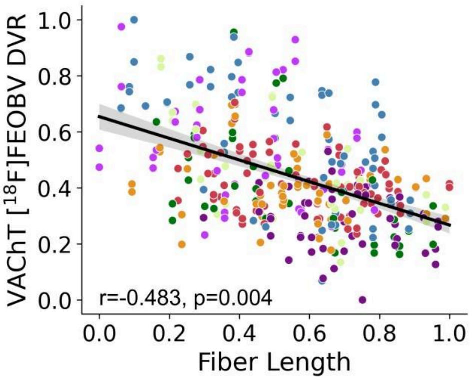


**Supplemental Fig 6. Correlation of basal forebrain (BF) white matter fiber lengths to cortical [$^{18}$F]FEOBV PET binding (VAChT).** Each point in the scatter plots represents cortical parcels based on HCP-MMP 1.0 parcellation color coded according to the Yeo networks[7] (Fig. 3B).

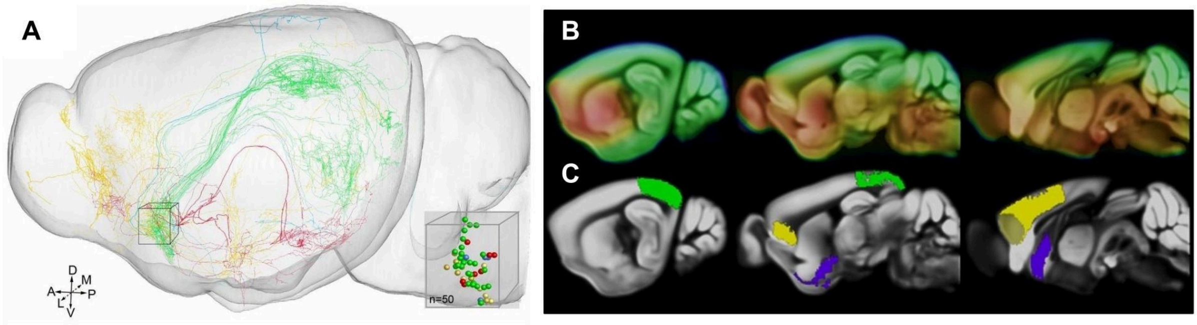


**Supplemental Fig 7. Cellular labeling of cholinergic neurons in the mouse brain.** (A) The half transparent sagittal view of the Allen Mouse Brain Atlas[8] is adapted from Li et al[9], showing axonal arborizations for 50 individual cholinergic neurons. The projections are color coded according to distinct anatomical locations they target. The soma of reconstructed neurons are shown In the 3D box inset with locations in medial septal nucleus and vertical band of Broca. (B) Average $[^{18}F]$FEOBV PET binding (Vt) in 11 wild type mice (VAChTflox/flox 108; C57BL/6J) at 6 months of age in Allen Mouse Brain Atlas common coordinate framework (CCF) v3 reference space (sagittal view). (C) The Allen Mouse Brain Atlas CCF v3 annotations for the visual cortex (green) and salience network regions (yellow), including cingulate, infralimbic and orbitofrontal cortices. For reference, the BF nuclei (medial septal nucleus, diagonal band of Broca and substantia innominata) are labeled in purple.
